# Dioecy in plants: an evolutionary dead end? Insights from a population genomics study in the *Silene* genus

**DOI:** 10.1101/414771

**Authors:** Aline Muyle, Hélène Martin, Niklaus Zemp, Maéva Mollion, Sophie Gallina, Raquel Tavares, Alexandre Silva, Thomas Bataillon, Alex Widmer, Sylvain Glémin, Pascal Touzet, Gabriel AB Marais

## Abstract

About 15,000 angiosperm species (∼6%) have separate sexes, a phenomenon known as dioecy. Early work reported a lower species richness in dioecious compared to non-dioecious sister clades, which was taken to suggest that dioecy might be an evolutionary dead end. More recently, phylogenetic analyses using different methodologies have challenged this conclusion. Here, we used a population genomics approach to look for evidence of evolutionary handicaps of dioecy in the *Silene* genus at the molecular level. We obtained RNA-seq data of individuals from several populations in 13 closely related species with different breeding systems: seven dioecious, three hermaphroditic and three gynodioecious species. We show that dioecy is associated with increased genetic diversity and a higher selection efficacy both against deleterious and for beneficial mutations while controlling for differences in population size. We conclude that, in the *Silene* genus, dioecious species bear no sign of mutational burden or upcoming extinction. On the contrary, dioecious species harbor a higher potential for adaptation than their non-dioecious relatives. Our results do not support the evolutionary dead end hypothesis and re-open the question why dioecy is rare in angiosperms.

**Significance statement:** Dioecy (=separate sexes) is much rarer in flowering plants compared to animals and other organisms. The “dead-end hypothesis” states that dioecious plant populations might experience evolutionary handicaps such as low seed dispersal (as only 50% of the individuals, the females, contribute), which might cause high genetic drift, low adaptation and ultimately extinction. Here we tested this hypothesis by focusing on the genus *Silene* and by comparing the population genetics of 13 dioecious and non-dioecious species. We found that dioecious *Silene* species exhibit lower genetic drift and more adaptation compared to their non-dioecious relatives. Our results thus reject the dead-end hypothesis and re-open the question of why dioecy is rare in flowering plants.

## Introduction

Out of 261,750 angiosperm species, approximately 15,600 (∼6%) are dioecious (1). Dioecy is thus rare compared to the dominant breeding system in angiosperms, hermaphroditism, in which individuals carry bisexual flowers. Nonetheless, dioecy can be found in ∼40% of angiosperm families. The current view is that many independent (871 to ∼5000) and mostly recent transitions occurred from hermaphroditism to dioecy in angiosperms (1). These events seem to have followed two main paths: one through a gynodioecious intermediate (with female and hermaphrodite individuals) called the gynodioecy pathway and another one through a monoecious intermediate (with individuals carrying both female and male flowers) called the monoecy or paradioecy pathway (2). In some cases, dioecy might have evolved through other minor pathways such as heterostyly (2).

Species richness was found to be lower in dioecious compared to non-dioecious sister clades, suggesting that dioecy might lead to an increased extinction rate in angiosperms (3). Some theoretical work explored this idea and showed that the hypothetically increased extinction rate could result from two main evolutionary handicaps. First, only females contribute to seed production and dispersal in dioecious species (4). Second, in dioecious species with insect pollination, male-male competition results in males being more attractive than females for pollinators, making reproduction and population size sensitive to variation in pollinator density (5). In angiosperms, dioecy is associated with a number of traits (fleshy fruits, wind pollination, inconspicuous flowers and woody growth) and ecological preferences (in the tropics and on islands). It has been proposed that handicaps of dioecy could drive these associations (6). For example, dioecy could be more frequent in species with wind pollination because these do not suffer from the “pollinator” handicap, which reduces their extinction risk.

The “evolutionary dead end” hypothesis has, however, been recently challenged (reviewed in 7). The sister clade analysis used in early work assumes that speciation events and trait transitions coincide (3). However, if dioecy evolved along one of the two branches formed by a speciation event, then equal species richness is not expected among sister clades even when diversification rates are similar (8). Using a corrected version of the sister clade analysis on an updated dataset, dioecious species were found to have higher species richness than non-dioecious sister clades in angiosperms, contrary to previous observation (9). Using a phylogenetic method to study trait evolution and species diversification (BiSSe, 10) on several genera with dioecious species, dioecy was not found to be associated with an increased extinction rate (11, 12). Another hypothesis has been put forward to explain the rareness of dioecy in angiosperms, which posits that reversion from dioecy to other breeding systems is frequent and that dioecy is more labile than previously thought (7). This is supported by a recent BiSSe analysis (13).

If dioecious species suffer from evolutionary handicaps, this should leave footprints in their genetic diversity. Seed-dispersal and pollinator handicaps should result in population size oscillations, which should reduce the long-term effective population size of a dioecious species. On the contrary, if dioecy is not a dead end but is instead beneficial in some ecological contexts, increased genetic diversity and faster adaptation should be observed in dioecious species (7). This question could be addressed through comparative population genetics of dioecious and related non-dioecious species, which has not been done yet (reviewed in 7).

In this work, we focused on 13 species of the large *Silene* genus (∼700 species) of the Caryophyllaceae family, in which dioecy has evolved at least twice (14). *Silene* is considered a model system to study the evolution of breeding systems in angiosperms, thanks to its well-described breeding systems and a reasonably robust phylogeny (15). Dioecy in *Silene* is likely to have evolved through the gynodioecious pathway (reviewed in 16). Dioecious *Silene* species should bear the hallmarks of evolutionary handicaps, if these exist, because they are short-lived herbs, many of them (but not all) are insect pollinated, they do not have fleshy fruits and they live in temperate regions. We sampled dioecious species from the two groups in which dioecy evolved independently (5 species in the *Melandrium* section and 2 species in the *Otites* section) along with several hermaphroditic and gynodioecious close relatives for each group and one outgroup (*Dianthus chinensis*). *Silene* species can have very large genome sizes (15) and no reference genome is available in the genus (only a fourth of the *Silene latifolia* genome has been assembled to date, 17). We therefore used a population genomics approach relying on RNA-seq data, which has proven successful in organisms that lack a reference genome (18–20). Multiple individuals were sampled across the species geographic ranges and sequenced by RNA-seq. A pipeline specifically designed for such data was applied to perform SNP-calling (18, 21, 22). The SNPs were then analyzed to estimate genetic diversity and the efficacy of selection in each species.

## Results and Discussion

### Phylogenetic and population genetics characterization of the sampled species

Our species sample (13 species in total, see Fig. 1) includes two hermaphroditic species (*S. viscosa* and *S. paradoxa*), three gynodioecious species (*S. vulgaris, S. nutans* W and *S. nutans* E) and seven dioecious species (section *Melandrium*: *S. heuffelii, S. dioica, S. marizii, S. diclinis, S. latifolia* and section *Otites*: *S. pseudotites* and *S. otites*) from subgenera *Behenantha* and *Silene*, the main subgenera in the *Silene* genus (23). For each species, we sampled from 4 to 34 individuals from different populations covering the geographic range of the species (see Fig. 1 for the number of individuals sampled in every species and Fig. S5 for sample locations). We sampled 130 individuals in total and sequenced them by RNA-seq (see Table S1 for details on samples). We then *de novo* assembled reference transcriptomes separately for each species and annotated ORFs and orthologs (see Material and Methods and Table S1). The dioecious *Silene* all have sex chromosomes and merging autosomal and sex-linked markers can be problematic in population genetics studies as they have different effective population sizes (e.g. 24). For downstream analysis, we thus only kept autosomal ORFs as predicted by the SEX-DETector method (25)(see Material and Methods).

**Fig. 1.**
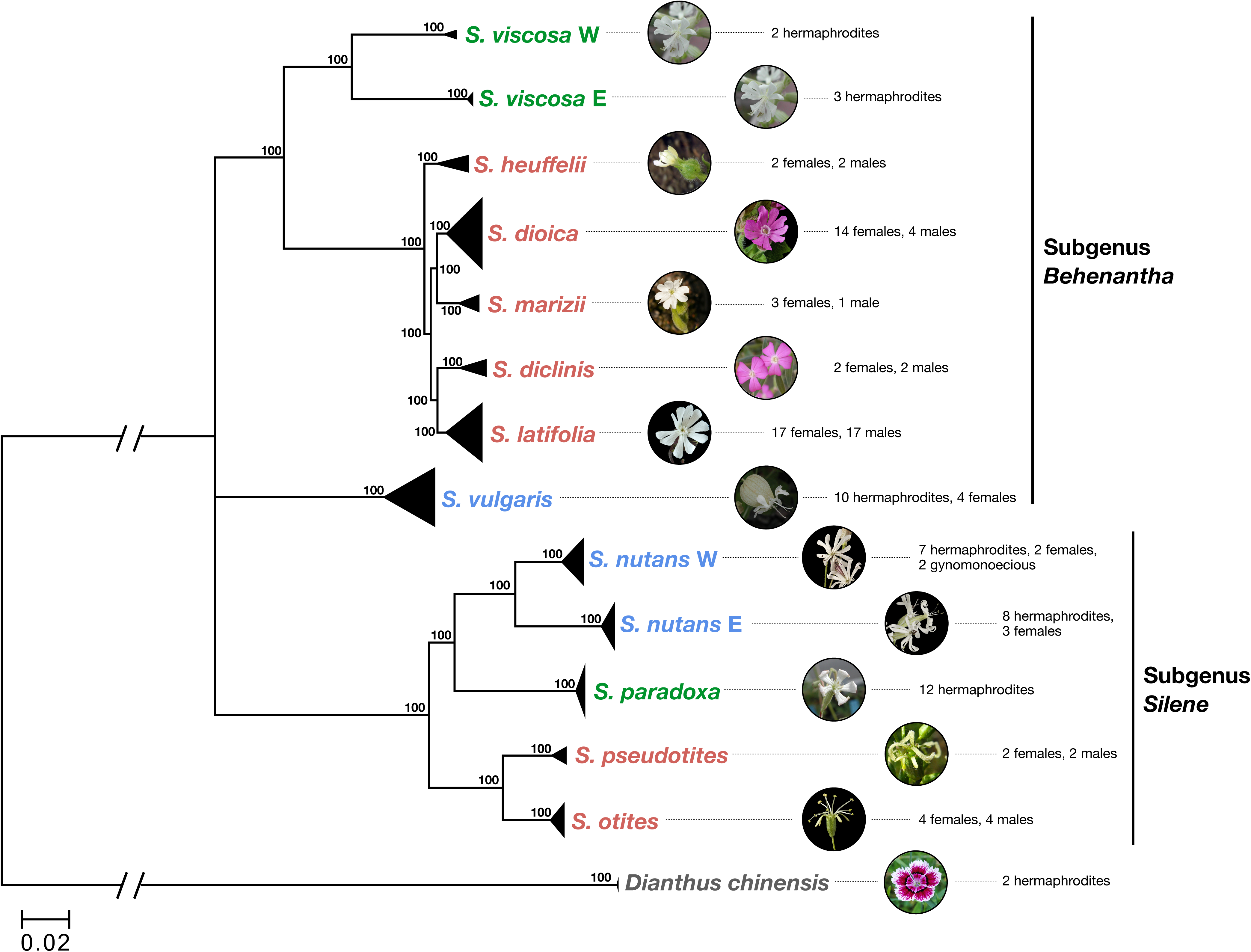
Sampled species and their phylogenetic relationships. Dioecious species are shown in red, gynodioecious species in blue and hermaphroditic species in green. Sample sizes and sex are indicated. Bootstrap support values were generated using 100 replicates. Pictures of *S. latifolia, S. dioica, S. nutans* W, *S. nutans* E, *S. otites* are from Maarten Strack van Schijndel, pictures of *S. marizii, S. vulgaris, S. paradoxa, S. viscosa* are from Oxelman *et al* (55), picture of *S. heuffelii* is from Dr Daniel L. Nickrent.

We built a phylogenetic tree to check for congruence with published phylogenies of the *Silene* genus (e.g. 23). Orthologous ORFs were aligned among *Silene* species and *Dianthus chinensis,* which was used to root the tree. A total of 55,337 SNPs spread over 1,294 contigs were used to build the tree (see Material and Methods). All samples from a given species (except *S. viscosa*) grouped together as expected (Fig. 1). The two lineages of *S. nutans* formed two clearly distinct groups that should be treated as distinct species as shown in previous studies (26, 27). *S. viscosa* samples formed two highly divergent lineages: a western lineage (*S. viscosa* W), which includes samples from Sweden and the Czech Republic, and an eastern lineage (*S. viscosa* E), which includes samples from Bulgaria, Russia and Kirgizstan. We treated these lineages as two differentiated taxa and analyzed them separately hereafter. Unlike in previous work (e.g., 28), *S. vulgaris* did not group with *S. viscosa* and the section *Melandrium*, but instead could not be reliably grouped with either subgenus, perhaps due to the limited number of species used to build the phylogeny here (Fig. 1).

We used an RNA-seq data specific pipeline to call SNPs (see Material and Methods). We found 1,088,269 SNPs in total (Table 1). We computed species-averaged heterozygosity levels and the Weir and Cockerham *F*_IT_ statistic, which indicates deviations from Hardy-Weinberg equilibrium such as selfing or population structure (see Material and Methods). In our set of species, the *F*_IT_ values are low to moderate and mainly reflect the effect of population structure as already observed (29), except for *S. viscosa* E that exhibits a high *F*_IT_ and very low heterozygosity and polymorphism levels, likely reflecting a high selfing rate (Fig. 2). For all other species, *F*_IT_ is equal to or lower than 0.2 so that population structure is not creating strong among-species difference in genetic diversity.

**Table 1.** Characteristics of the datasets and coding sequence synonymous polymorphism (π_S_), non- synonymous (π_N_) for all studied species.

| Breeding system | Focal species | Number of samples | Number of clean contigs | Number of snps | Weir & Cockerham $F_{IT}$ | $\pi_S$ in focal species (%) | $\pi_N$ in focal species (%) | $\pi_N/\pi_S$ |
| --- | --- | --- | --- | --- | --- | --- | --- | --- |
| **Section *Benantha*** |  |  |  |  |  |  |  |  |
| dioecious | *S. diclinis* | 3 | 2729 | 23774 | -0.089<br>[-0.113;-0.065] | 0.934<br>[0.886;0.982] | 0.179<br>[0.170;0.187] | 0.191<br>[0.182;0.200] |
|  | *S. dioica* | 17 | 3730 | 143297 | 0.181<br>[0.176;0.186] | 2.073<br>[2.013;2.135] | 0.242<br>[0.234;0.251] | 0.117<br>[0.113;0.121] |
|  | *S. heuffelli* | 4 | 3999 | 98957 | -0.043<br>[-0.051;-0.035] | 2.698<br>[2.628;2.771] | 0.342<br>[0.331;0.353] | 0.127<br>[0.123;0.131] |
|  | *S. latifolia* | 34 | 4132 | 214646 | 0.165<br>[0.161;0.169] | 2.005<br>[1.950;2.062] | 0.244<br>[0.236;0.252] | 0.122<br>[0.118;0.126] |
|  | *S. marizii* | 4 | 2878 | 44868 | 0.013<br>[-0.005;0.030] | 1.637<br>[1.573;1.703] | 0.246<br>[0.235;0.256] | 0.150<br>[0.144;0.156] |
| gynodioecious | *S. vulgaris* | 14 | 6248 | 281300 | 0.205<br>[0.200;0.209] | 2.685<br>[2.639;2.732] | 0.351<br>[0.343;0.360] | 0.131<br>[0.128;0.134] |
| hermaphrodite | *S. viscosa* E | 3 | 6413 | 13712 | 0.723<br>[0.691;0.754] | 0.291<br>[0.277;0.305] | 0.056<br>[0.053;0.059] | 0.194<br>[0.184;0.204] |
|  | *S. viscosa* W | 2 | 5959 | 24305 | 0.068<br>[0.038;0.098] | 0.723<br>[0.693;0.757] | 0.134<br>[0.127;0.142] | 0.186<br>[0.178;0.193] |
| **Section *Otites*** |  |  |  |  |  |  |  |  |
| dioecious | *S. pseudotites* | 4 | 2339 | 24698 | 0.024<br>[0.006;0.041] | 1.116<br>[1.074;1.157] | 0.195<br>[0.185;0.204] | 0.174<br>[0.166;0.183] |
|  | *S. otites* | 8 | 893 | 11364 | 0.014<br>[-0.007;0.035] | 1.081<br>[1.016;1.150] | 0.180<br>[0.167;0.194] | 0.167<br>[0.154;0.180] |
| gynodioecious | *S. nutans* E | 11 | 5100 | 65820 | 0.124<br>[0.106;0.140] | 0.861<br>[0.829;0.895] | 0.130<br>[0.125;0.136] | 0.151<br>[0.146;0.157] |
|  | *S. nutans* W | 11 | 5059 | 88853 | 0.201<br>[0.182;0.219] | 1.048<br>[1.011;1.084] | 0.170<br>[0.165;0.176] | 0.163<br>[0.157;0.169] |
| hermaphrodite | *S. paradoxa* | 12 | 5955 | 52675 | 0.116<br>[0.108;0.124] | 0.589<br>[0.568;0.611] | 0.102<br>[0.098;0.105] | 0.172<br>[0.166;0.179] |

**Fig. 2.**
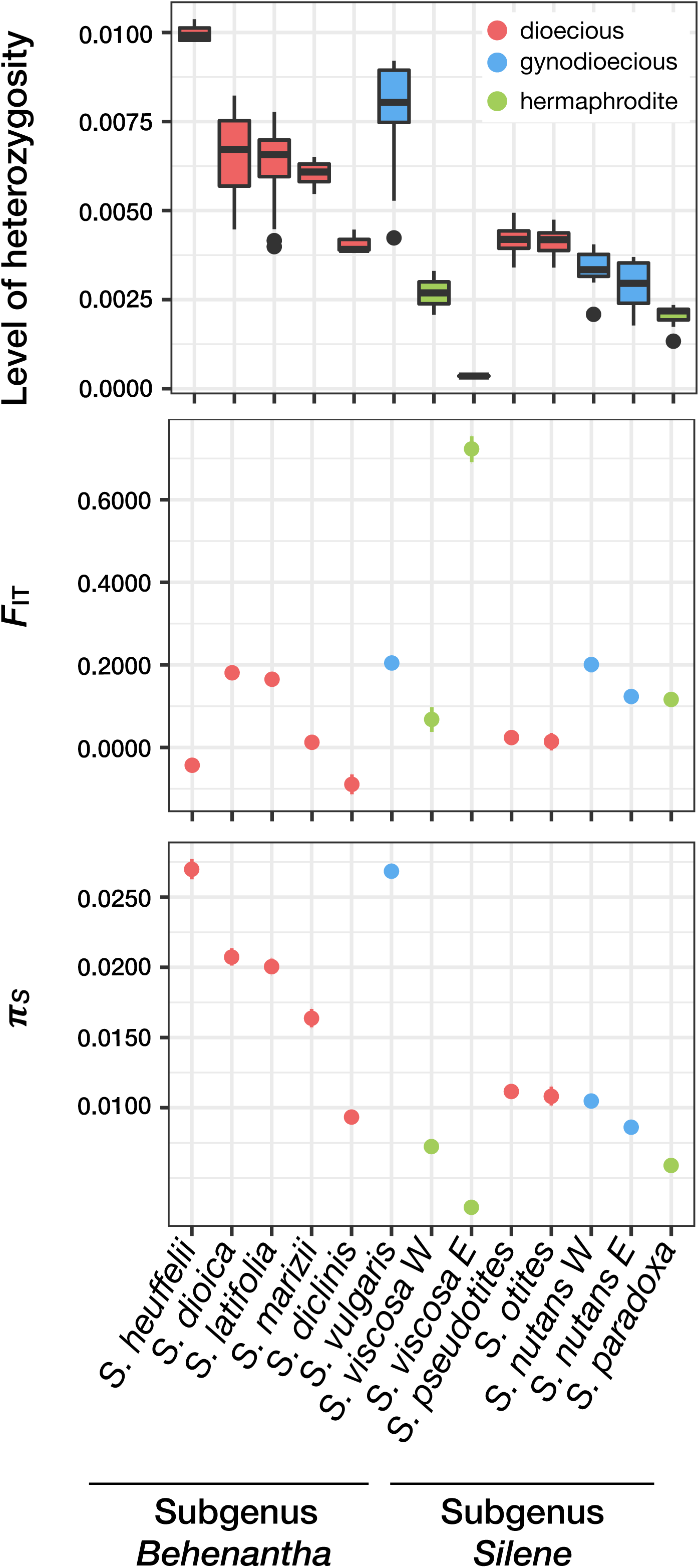
Levels of individual heterozygosity (proportion of heterozygote positions), Weir and Cockerham *F*_IT_ statistic and π_S_ within each species. Dioecious species are shown in red, gynodioecious species are shown in blue and hermaphroditic species are shown in green.

### Genetic diversity is higher in dioecious *Silene* species

We tested the evolutionary dead-end hypothesis by computing the synonymous genetic diversity (π_S_), which reflects the effective population size (*N*_*e*_) with π_S_*=4N*_*e*_*μ* where *μ* is the mutation rate (see Material and Methods). *N*_*e*_ is a direct measure of the intensity of drift in populations: under the evolutionary dead-end hypothesis, and assuming the mutation rate is constant among species, we expect lower values of π_S_ for dioecious compared to non-dioecious species. Table 1 and Fig. 2 show the opposite trend: dioecious species have higher π_S_ values than non-dioecious species.

The 13 species have very different census population sizes (*N*) with some species being cosmopolitan (e.g. *S. latifolia, S. vulgaris*) and others rare and endemic (e.g. *S. diclinis*). The genetic diversity is somewhat affected by *N*. Although this relationship can be complex (as *N* and *N*_*e*_ can be quite different), we need to control for the effect of *N* to extract the specific effect of breeding systems on genetic diversity. Our proxies for *N* were (i) the number of occurrences of each species in the Global Biodiversity Information Facility (GBIF) dataset, and (ii) the geographic range of each species according to the Atlas *Florae Europaeae* (see Material and Methods), both of which should correlate with *N*.

We used a linear model to jointly assess the effects of *N* and the breeding system on π_S_. GBIF data and Atlas geographic range (estimates of *N*) have a significant positive effect on π_S_ (*p*-values =0.030 and 0.012, t-values = 2.529 and 3.054, R^2^ = 0.390 and 0.483 respectively, and see Fig. 3). This shows that genetic diversity is strongly influenced by *N* in our set of *Silene* species. After controlling for the effect of *N* on genetic diversity, we found that dioecious species had a significantly higher π_S_ than non-dioecious species (linear model pairwise t-test *p*-values = 0.142 and 0.048, t-values = 1.595 and 2.247 using GBIF data and Atlas geographic range estimates respectively). In this analysis, different sets of genes were used in different species. As this might affect the results, we repeated the analysis on two smaller sets of genes, one shared by every species in the subgenus *Behenantha* and the second shared by every species in the subgenus *Silene*. Similar results were found using these shared gene subsets, i.e. GBIF data and Atlas geographic range have a significant positive effect on π_S_ (*p*-values = 0.021 and 0.012, t-values = 2.725 and 3.074, R^2^ =0.429 and 0.488, respectively). Dioecious species had higher π_S_ compared to other species after correcting for *N* (linear model pairwise t-test *p*-values = 0.127 and 0.045, t-values = 1.662 and2.283 using GBIF data and Atlas geographic range respectively, see Fig. S1).

**Fig. 3.**
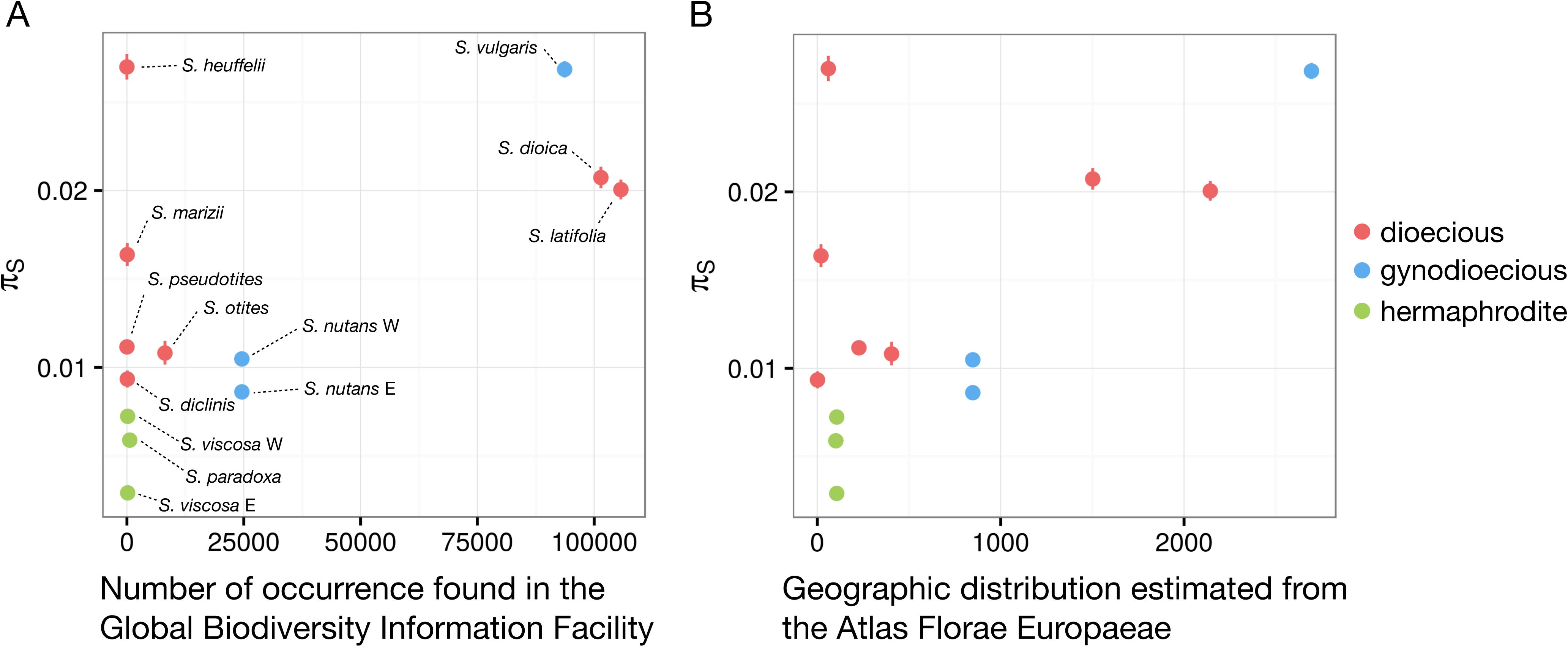
Relationship between synonymous polymorphism (π_S_) and **(A)** census population size estimated from the Global Biodiversity Information Facility (GBIF) and **(B)** geographic range estimated from the Atlas *Florae Europaeae*. For each species, dots represent the average π_S_ and lines mark the confidence interval of π_S_. Census population sizes estimated from GBIF refer to the number of occurrences, and geographic distributions from the Atlas *Florae Europaeae* refer to the number of dots (native occurrence) counted on the Atlas maps (see Material and Methods). Dioecious species are shown in red, gynodioecious species are shown in blue and hermaphroditic species are shown in green.

### Selection is more efficient in dioecious *Silene* species

We then tested whether selection was weaker in dioecious species as predicted by the evolutionary dead-end hypothesis. We did not use the ratio of non-synonymous to synonymous substitution rate d_N_/d_S_as we are studying closely related species and some shifts in breeding systems might have occurred very recently (14, 15). We rather looked at the ratio of non-synonymous over synonymous polymorphism π_N_/π_S_, a more appropriate statistic to study the efficacy of selection on a short evolutionary timescale (30). Increased π_N_/π_S_ values are expected when purifying selection is reduced. We found no statistical association between breeding systems and π_N_/π_S_ (Table 1). Interpreting patterns of π_N_/π_S_ is, however, not trivial when mutations are affected by both purifying and positive selection.

We thus turned to a more sophisticated approach to study selection and applied a modified version of the McDonald-Kreitman framework called polyDFE to our data (31). This framework estimates the distribution of fitness effects (DFE) of both deleterious and beneficial mutations using polymorphism data only, thus avoiding the problem of recent shifts in population sizes when both polymorphism and divergence data is used, as in other frameworks (32, 33). Note that we could not include *S. viscosa* E in the analysis: as it is highly selfing only one allele per individual should have been sampled, leading to a too small site frequency spectrum for proper inferences (*n* = 3) (see Material and Methods). PolyDFE estimated the DFE (Fig. S2) for each species with a gamma distribution for deleterious mutations and exponential distribution for beneficial mutations (see estimated parameter ranges in Fig. S3). From this, we obtained the predicted proportion of adaptive substitutions (α) and the predicted rates of adaptive substitutions (ω_A_) and of deleterious substitutions (ω_NA_) (31). Under the dead-end hypothesis, we expect dioecious species to have a higher ω_NA_ and a lower α and ω_A_ because of a faster accumulation of deleterious mutations and a lower rate of adaptation compared to non-dioecious species.

We used a linear model to jointly assess the effects of *N*_*e*_ (using π_S_ as a proxy) and of the breeding system on α. We found a significant positive effect of π_S_ on α (*p*-value = 0.003, t-value = 4.07, Fig. 4). This has previously also been observed in animal species (32) where the correlation between α and π_S_ was solely due to the link between π_S_ and ω_NA_. Because α = ω_A_ / (ω_A_ + ω_NA_), when selection is more efficient (higher *N*_*e*_ and higher π_S_), the rate of deleterious substitutions ω_NA_ is reduced, which in turn increases α. However, in our dataset, the correlation between α and π_S_ was both due to higher adaptive evolutionary rates and lower deleterious evolutionary rates when effective population size is high. Indeed, π_S_ had a significant positive effect on ω_A_ (linear model *p*-value = 0.011, t-value = 3.19) and a significant negative effect on ω_NA_ (linear model p-value = 0.00017, t- value = −6.15, see Fig. 4).

**Fig. 4.**
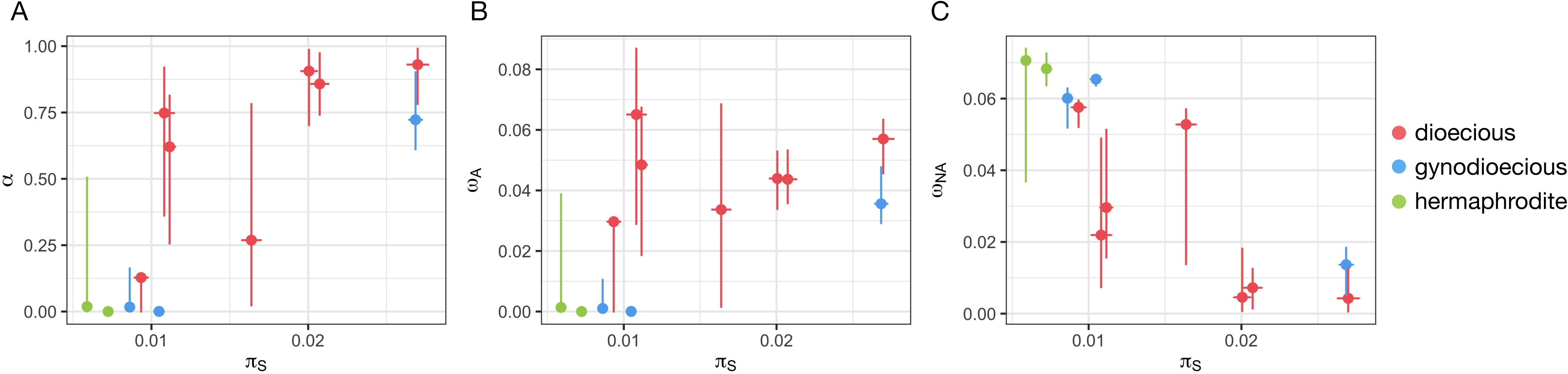
Relationship between synonymous polymorphisms (π_S_) and **(A)** the proportion of adaptive amino-acid substitutions, α, **(B)** the rate of adaptive non-synonymous substitutions ω_A_, **(C)** the rate of non-adaptive non-synonymous substitutions, ω_NA_. Dioecious species are shown in red, gynodioecious species are shown in blue and hermaphroditic species are shown in green.

We then tested the dead-end hypothesis by comparing α, ω_A_ and ω_NA_ among dioecious and non-dioecious species, after controlling for the effect of *N*_*e*_. We found that non-dioecious species had significantly lower α and ω_A_ and a significantly higher ω_NA_ compared to dioecious species (linear model pairwise t-test *p*-values = 0.015, 0.000311 and 0.018547, t-values = −2.998, −5.657 and 2.868 for α, ω_A_ and ω_NA_ respectively). Dioecious species thus experience fewer fixations of slightly deleterious mutations due to genetic drift and more adaptive evolution, which is the opposite of the expected trend under the dead-end hypothesis. We repeated the analysis using orthologous genes shared within the subgenus *Behenantha* or within the subgenus *Silene* as different sets of genes in different species could affect the results. The conclusions, however, remained unchanged, although statistical power was reduced and some differences were no longer significant (see Fig. S4 legend for details).

### Concluding remarks

Overall, our findings show that dioecious *Silene* species exhibit higher genetic diversity, experience more efficient purifying selection and benefit from increased adaptive evolution compared to their non-dioecious relatives. This is clearly not supporting the dead-end hypothesis, and suggests on the contrary that gaining dioecy is beneficial in the *Silene* genus. What dioecious *Silene* species are adapting to, however, remains to be understood. Obligate outcrossing could improve adaptation of colonizing species such as *S. latifolia* and *S. dioica*, but would not explain well why higher rates of adaptation are also found in endemic species such as *S. diclinis*. Sexual conflicts, sexual selection and sexual dimorphism, which are typical of dioecious species could drive the patterns of positive selection that we observed in *Silene*. Such selection was shown to affect gene expression and result in the evolution of many sex-biased genes in *S. latifolia* (34). Coding regions could be affected similarly but we excluded potentially sex-linked genes, removing the main target of these forces. Reproductive assurance provides an ecological advantage to hermaphroditic self-compatible species over dioecious species as few or even a single individual can start a new population (35). The other side of the coin is that hermaphroditic self-compatible species can experience recurrent bottlenecks strongly reducing genetic diversity and rendering selection less efficient. On the contrary, because of the so-called Allee effect (i.e. lower population growth rate at low densities), dioecious populations cannot decline below a given threshold at risk of declining to extinction, which can paradoxically help maintaining high genetic diversity in populations that have persisted so far (36). Here we focused on polymorphism data, which give a snapshot of the evolutionary dynamics of a species. Looking at these dynamics over longer evolutionary timescales would be interesting. Käfer et al. (37) compared patterns of divergence between dioecious and non-dioecious *Silene*. A higher d_N_/d_S_, suggestive of a stronger genetic drift, was found in dioecious compared to non-dioecious species in the *Behenantha*, but not in the *Silene* subgenera. Whether this means that dioecy has not always been beneficial in *Silene* or whether this is merely a consequence of their dataset being small remains to be addressed with more data. If dioecy is as beneficial as suggested here, why would dioecy be so rare in angiosperms? Of course, obligate outcrossing can be achieved through different mechanisms in angiosperms, not only dioecy (2). Moreover, dioecy might be much more evolutionary labile than previously thought, being easily gained and lost depending on environmental conditions (7). Our work re-opens the fascinating question of the rareness of dioecy in angiosperms.

## Material and Methods

### Sampling and sequencing

We investigated patterns of polymorphism and divergence of three breeding systems (hermaphroditism, gynodioecy and dioecy) in two subgenera of the *Silene* genus, subgenus *Behenantha* and subgenus *Silene* (sample size and sexes are indicated in Fig. 1). For the subgenus *Behenantha*, we sampled five dioecious species (*S. latifolia, S. diclinis, S. dioica, S. marizii* and *S. heuffelli*), one gynodioecious species (*S. vulgaris*), and one hermaphroditic species (*S. viscosa*). For the subgenus *Silene*, we sampled two dioecious species (*S. otites* and *S. pseudotites*), two gynodioecious sub-species (*S. nutans* W - Western lineage, and *S. nutans* E - Eastern lineage, as defined in 26) and one hermaphroditic species (*S. paradoxa*). Two samples of *Dianthus chinensis* were added to root the phylogeny. Some of the individuals were already used in previous studies (S1).

Seeds were sampled across the geographic distribution of each species (Fig. S5) and were sown and grown in controlled greenhouse conditions. In case plants did not flower, young leaves were sampled instead (details on individual names, sex, origin and sampled tissue can be found in Table S1). Total RNA was extracted through the Spectrum Plant Total RNA kit (Sigma, Inc., USA) following the manufacturer’s protocol and treated with DNAse. Libraries were prepared with the TruSeq RNA sample Preparation v2 kit (Illumina Inc., USA). Each 2 nM cDNA library was sequenced using a paired-end protocol on an Illumina HiSeq2000 sequencer, producing 100 bp reads. Demultiplexing was performed using CASAVA 1.8.1 (Illumina) to produce paired-sequence files containing reads for each sample in Illumina FASTQ format. RNA extraction and sequencing were done by the sequencing platform in the AGAP laboratory, Montpellier, France (http://umr-agap.cirad.fr/). The Melandrium and *Dianthus* samples were sequenced as described in Zemp et al. (34) at the Quantitative Genomics Facility (QGF, ETH Zürich, Switzerland) and by GATC Biotech AG (Konstanz, Germany).

### Reference transcriptome assemblies

Reference transcriptomes were assembled *de novo* for all species (except for *S. marizii, S. heuffelli, S. diclinis* and *S. dioica*). For the subgenus *Behenantha*, reads from all individuals were pooled for each species. Steps in the assembly are represented in Fig. S3. 100% identical reads, assumed to be PCR duplicates, were filtered using the ConDeTri trimming software (38). Illumina reads were then filtered for sequencing adapters and low read quality using the ea-utils FASTQ processing utilities (39). Filtered reads were assembled using trinity with default settings (40). Poly-A tails were removed using PRINSEQ (41) with parameters -trim_tail_left 5 -trim_tail_right 5. rRNA-like sequences were removed using riboPicker version 0.4.3 (42) with parameters -i 90 -c 50 -l 50 and the following databases: SILVA Large subunit reference database, SILVA Small subunit reference database, the GreenGenes database and the Rfam database. Obtained transcripts were further assembled, inside of trinity components, using the program Cap3 (43) with parameter -p 90 and home-made perl scripts.

Raw data of the five taxa of subgenus *Silene* were filtered for sequencing adapters using Cutadapt (44), low read quality and poly-A tails with PRINSEQ (41). For each focal species of the subegnus, reference transcriptomes were assembled *de novo* mixing filtered reads from two individuals from the same population using default settings of Trinity (40) and an additional Cap3 run (43). Finally, in both subgenus, coding sequences were predicted using TransDecoder (see website: http://transdecoder.github.io/, 40) with PFAM domain searches as ORF retention criteria. Assembly statistics can be found in Table S2.

### Mapping, genotyping and alignment

Reads from each species were mapped onto their respective reference transcriptome using BWA version 0.7.12 with parameter -n 5 (45) for the subgenus *Behenantha* and Bowtie2 (46) for the subgenus *Silene*. Mapping statistics can be found in Table S3. For the *Melandrium* section (*S. latifolia, S. marizii, S. heuffelli, S. diclinis* and *S. dioica*), all reads were mapped to the *S. latifolia* reference transcriptome presented above. SAM files were compressed and sorted using SAMtools (47). Individuals were genotyped using reads2snp_2.0 (18, 21) with parameters -par 1 -min 3 (10 for the subgenus *Silene*) -aeb -bqt 20 -rqt 10 that is to say: stringent filtering of paralogous positions, minimum coverage of 3 (or 10) reads for calling a genotype, accounting for allelic expression bias, minimum read mapping quality threshold of 10 and minimum base quality threshold of 20. The output obtained was two unphased sequences for each individual and each ORFs.

Orthologous ORFs were identified using OrthoMCL Basic Protocol 2 (48) with default parameters. Best reciprocal blast hits were considered orthologs among species and orthologous ORFS were aligned for all individuals using MACSE (49). Species from the subgenus *Behenantha* were aligned separately with *S. paradoxa*, considered as an outgroup of this section and species from the subgenus *Silene* were aligned with *S. viscosa* considered as an outgroup of this section.

### Phylogenetic reconstruction

Orthologous ORFs among species were obtained using best reciprocal blast hits among the nine species (all species from the *Silene* genus and *Dianthus chinensis*). For each orthologous ORF, the nine species were added one by one into the alignment using MACSE (49). The data were filtered so that a maximum of 20% missing data per site and per species was allowed, for a total of 47,075 SNPs spread over 1,345 ORFs. We used the maximum number of ORFs available and did not restrict our analysis to autosomal ORFs identified in dioecious species. The phylogenetic tree was reconstructed using the GTRGAMMA model in RaxML (50). Bootstrap support values were generated using 100 replications.

### Data filtering

Analyses were carried on autosomal genes only, in order to avoid potential confounding effects of sex chromosomes and breeding systems on selection efficiency. Autosomal genes were predicted by SEX-DETector (25) thanks to the study of a cross (parents and progeny) sequenced by RNA-seq (Martin et al. *submitted* and Muyle et al. *in prep*). 11,951 contigs were inferred as autosomal in *S. latifolia*, 7,695 for *S. diclinis*, 11,675 for *S. heuffelli*, 8,997 for *S. marizii*, 10,698 for *S. dioica*, 2,035 in *S. otites* and 6,234 in *S. pseudotites*.

The data were further filtered in order to remove ORFs with frameshifts. Only biallelic sites were kept for analyses (sites with two or less alleles). Only ORFs longer than 100 codons and with a maximum of 20% of missing individuals of the focal species in the alignment were retained, after removing incomplete codons and gaps. We obtained final datasets from 893 retained ORFs for *S. otites* to 6,413 retained ORFs for *S. viscosa* E (see Table 1). Finally, to verify that observed trends are not due to differences in studied ORFs among species, we performed the same analyses on a smaller set of orthologous ORFs present in all species within a subgenus, so that parameters were estimated on the same set of ORFs. For these tests, 188 orthologous ORFs were analysed for the subgenus *Silene*, and 298 for the subgenus *Behenantha*.

### Polymorphism, heterozygosity and F_IT_

Population statistics were computed on the filtered dataset for each ORF and averaged using the PopPhyl tool dNdSpiNpiS_1.0 (18) with parameter kappa=2. The following statistics were computed for each contig: the mean fixation index *F*_IT_ (51), per-individual heterozygosity (proportion of heterozygote positions), per-site synonymous polymorphism (π_S_), per-site non-synonymous polymorphism (π_N_) and the ratio of non-synonymous to synonymous polymorphism (π_N_/π_S_). Note that we could not compute *F*_IS_ or *F*_ST_ as we do not have the required 2 individuals / population. Confidence intervals were obtained by 1,000 bootstraps on contigs.

### Estimation of geographic distribution and population size

Geographic distributions of the *Silene* species were estimated by counting the black dots (native occurrence) on the Atlas maps by Jalas and Suominen (52). This was partly achieved using the Dot Count program (http://reuter.mit.edu/software/dotcount). Since no map was available for *S. pseudotites*, dots were counted for Italy and adjacent regions of France and Ex-Yougoslavia using the *S. vulgaris* map (the Atlas indicates these are the regions where this plant is abundant).

Census population sizes of the *Silene* species were estimated using raw species occurrence numbers from GBIF (http://www.gbif.org 17th January 2017 Download of *S. dioica*: http://doi.org/10.15468/dl.fim8f2, *S. marizii*: http://doi.org/10.15468/dl.yywbrk, *S. diclinis*: http://doi.org/10.15468/dl.vizuiv and 1st December 2016 Download of *S. pseudotites*: http://doi.org/10.15468/dl.yrvenq, *S. otites*: http://doi.org/10.15468/dl.fxoocf, *S. paradoxa*: http://doi.org/10.15468/dl.wizsb6, *S. nutans*: http://doi.org/10.15468/dl.poqxc1, *S. viscosa*: http://doi.org/10.15468/dl.bwdu9z, *S. vulgaris*: http://doi.org/10.15468/dl.vmvebr, *S. latifolia*: http://doi.org/10.15468/dl.mfvyds). Some countries contribute poorly to this database, however, we assumed that this affected all *Silene* species in the same way. The GBIF database was filtered for showing occurrence from human observation and literature occurrence. Only data from European countries and Russia, which represent the native range of those species, were kept. For each species, all existing names of the species were included.

Both estimates (geographic range and population size) were divided by two when two subspecies were identified with our phylogeny (i.e. for *S. viscosa* and *S. nutans*). Geographic range and GBIF data were correlated to π_S_ and breeding systems using R (53) linear model function lm as follows: lm(piS ∼ Atlas + mating_system)

lm(piS ∼ GBIF + mating_system)

### Selection efficacy

The unfolded synonymous and non-synonymous site frequency spectra (SFS, the distribution of derived allele counts across SNPs) were computed for each species with the sfs program provided by Nicolas Galtier (32). These SFS were used to estimate α (the proportion of adaptive amino-acid substitutions) in each species. Estimation of α was made according to a modified version of the Eyre-Walker and Keightley method (33), that relies on polymorphism data only to estimate α (31). The program was used to estimate the distribution of fitness effects of mutations (DFEM), α, the (per synonymous substitution) rate of adaptive non-synonymous substitution ω_A_=α d_N_/d_S_ and the rate of non-adaptive non-synonymous substitutions ω_NA_=(1-α) d_N_/d_S_. For each species separately, four different models were fitted to the SFS:

- Model 1: full DFEM with adaptive mutations (p_b > 0), estimation of an orientation error rate for ancestral alleles in the SFS (eps_an), estimation of demography (r_i ≠ 1) for a total of 6 parameters plus the number of demography parameters r_i which depends on the sample size of each species.
- Model 2: same as Model 1 but without ancestral allele orientation error rate (eps_an=0), for a total of 5 parameters plus the number of demography parameters r_i.
- Model 3: same as Model 1 but without adaptive mutations (p_b=0), for a total of 5 parameters plus the number of demography parameters r_i.
- Model 4: same as Model 1 but without adaptive mutations (p_b=0) and without ancestral allele orientation error rate (eps_an=0), for a total of 4 parameters plus the number of demography parameters r_i.

Each model provided estimates for α, ω_A_ and ω_NA_. These values were averaged using Akaike weigths as follows:

where *m* stands for model number *m, α*_*m*_ stands for the estimated α using model *m*, and

with AIC the Akaike criterion:

Where *n*_*m*_ stands for the number of parameters and *L*_*m*_ the likelihood of model *m*. Average values among models for ω_A_ and ω_NA_ were obtained using a similar procedure.

For each species, 500 bootstrapped datasets were randomly generated by random sampling SNPs with replacement from the original SFS. Average values among models were computed as explained previously for each bootstrap. The distribution of values for the 500 bootstraps was used to compute approximate 95% confidence intervals for α, ω_A_ and ω_NA_. For this, 2.5% of values were removed from each side of the bootstrap distribution (54). α, ωA and ωNA values were analysed in a linear model using R (lm function, 53). The model tested the effect of π_S_ and breeding systems (dioecious, gynodioecious, hermaphroditic) as fixed effect, while taking into account the variance of the estimations with weights computed as follow:

lm(α ∼ π_S_ + breeding system, weights) lm(ω_A_ ∼ π_S_ + breeding system, weights) lm(ω_NA_ ∼ π_S_ + breeding system, weights)

## Acknowledgements

We would like to thank Jos Kafer for his help with fieldwork and a number of colleagues for sending us seeds: J. J. Wieringa, Deborah Charlesworth, Bohuslav Janousek, Tatiana Giraud, Honor C. Prentice, Salvatore Cozzolino, Delphine Griveau, Henk Schat, Cristina Gonnelli, Alessia Guggisberger, Sophie Karrenberg, Adrien Favre, Maria Domenica Moccia, Aria Minder, Carmen, Max and Kerstin Kustas. AM, RT and GABM thank Elise Lacroix technician of the Univ. Lyon 1 greenhouse and and HM, SG and PT thank Eric Schmitt for taking care of the *Silene* plants. We are grateful to Sylvain Santoni and the AGAP platform staff for the RNA extraction and RNAseq. The numerical results presented in this paper were carried out using the computing cluster of Laboratoire Biométrie et Biologie-Pôle Rhone-Alpin de Bioinformatique (LBBE-PRABI) at Univ. Lyon 1 and the HPC service of the Centre de Ressources Informatiques (CRI) of Univ. Lille 1, for which we thank Stéphane Delmotte and Bruno Spataro. We thank Nicolas Galtier for help with his program sfs. HM, SG and PT are grateful to Jonathan Aceituno for technical support in R and to Céline Poux for discussion on phylogenetic inference. This work was supported by the Agence Nationale de la Recherche (ANR-11-BSV7-013-03, TRANS) to SG, GM, and PT, the Swiss National Science Foundation (31003A-182675), and a PhD fellowship from the French Research Ministry to HM.

